# Recovery from social isolation requires dopamine in males, but not the autism-related gene *nlg3* in either sex

**DOI:** 10.1101/2023.12.05.570093

**Authors:** Ryley T. Yost, Andrew M. Scott, Judy M. Kurbaj, Brendan Walshe-Roussel, Reuven Dukas, Anne F. Simon

## Abstract

Social isolation causes profound changes in social behaviour in a variety of species including humans, monkeys, mice, bees, and vinegar flies. However, the genetic and molecular mechanisms modulating behavioural responses to both social isolation and social recovery remain to be elucidated. In this study, we quantified the behavioural response of vinegar flies to social isolation through the use of two distinct protocols, one involving flies’ social space preference and the other assessing flies’ sociability, defined as their spontaneous tendencies to form groups. We found that social isolation increased social space and reduced sociability. These effects of social isolation, however, were reversible and could be reduced after 3 days of group housing. Flies with a loss of function of *neuroligin3* (ortholog of autism-related *neuroligin* genes) with known increased social space in a socially enriched environment, were still able to recover from social isolation. Using a *UAS-TH*-RNAi driven in all neurons, we show that dopamine is important for a response to social isolation and recovery in males but not in females. Furthermore, only in males, dopamine levels are reduced after isolation and are not recovered after group housing. Finally, in socially enriched flies with a loss of function of *neuroligin3*, dopamine levels are reduced in males, but not in females. We propose a model to explain how dopamine and *neuroligin3* are involved in the behavioural response to social isolation and its recovery in a dynamic and sex-specific manner.

## 1 Introduction

The environment an individual has been exposed to can influence social behaviour by affecting the interactions between individuals (Rubenstein and Hofmann, 2015). Social isolation, or the absence of interactions with others, has profound effects on mental and physical health in humans (Holt-Lunstad et al., 2010; Cacioppo et al., 2011; Meyer-Lindenberg and Tost, 2012). During the COVID-19 times, forced social isolation had negative consequences on families and individuals (Pellicano and Stears, 2020). Higher risk individuals such as children and people with neuropsychiatric disorders, like Autism Spectrum Disorders, face more challenges to their physical and mental health because of changes to their behavioural and environment support (Bellomo et al., 2020; Summers et al., 2021). It is important to understand the interactions between genetic predispositions and altered social environment, as well as the possibility to recover from social isolation.

Social isolation causes changes to the social behaviour in several organisms, including a variety of behavioural deficits in monkeys (Harlow et al., 1965;Mitchell and Clark, 1968;Harlow and Suomi, 1971;Suomi et al., 2008), deficits in social interactions, aggression, fear and anxiety in mice (Wongwitdecha and Marsden, 1996; Heidbreder et al., 2000; Fone and Porkess, 2008; Zhang et al., 2012; An et al., 2017; Endo et al., 2018; Medendorp et al., 2018; Zhang et al., 2018) and decreased social affiliation in bees (Hewlett *et al*., 2018). In *Drosophila melanogaster*, social isolation leads to physiological and behavioural changes, including decreased fiber number in the mushroom bodies (Technau, 1984), changes to neural excitability (Ueda and Wu, 2009), decreased lifespan (Leech *et al*., 2017), chemical communication (Kent *et al*., 2008), olfactory memory (Chabaud *et al*., 2009), courtship and courtship memory (Krupp *et al*., 2008), sleep patterns (Ganguly-Fitzgerald *et al*., 2006), circadian rhythm (Krupp *et al*., 2008), aggression (Ueda and Kidokoro, 2002; Ueda and Wu, 2009;Ramin et al., 2014), sociability (Scott et al., 2018) and social network structure (Bentzur *et al*., 2020). Isolated male flies are much more territorial and defend food patches and access to mates more than socially enriched flies (Hoffman, 1990). Our work and others’ have identified that isolated flies have increased social space (Simon et al., 2012; Xie et al., 2018; Yost et al., 2020), a measure of inter-individual distances (Simon et al., 2012; Burg et al., 2013).

Plastic changes in behaviour are important for an organism to adapt to a changing environment. Changes in social behaviour because of isolation can be partially or fully recovered in monkeys (Harlow et al., 1965; Harlow and Suomi, 1971), mice (Chen et al., 2016; An et al., 2017; Endo et al., 2018) and bees (Hewlett *et al*., 2018). Flies are no exception. In *Drosophila*, sleep and aggression levels were recovered after group housing (Ganguly-Fitzgerald et al., 2006; Wang et al., 2008). However, more research needs to be conducted on the recovery from isolation of other *Drosophila* social behaviours.

Previous studies have identified sensory modalities, neurocircuitry and synaptic proteins to be important for social space (Brenman-Suttner *et al*., 2018). One identified synaptic protein is *neuroligin3*, a post-synaptic cell adhesion molecule that regulates synaptic development and function and is an ortholog to a human autism candidate gene (Knight et al., 2011; Sindi et al., 2014; Yost et al., 2020). We have previously shown *nlg3* to be important for social space (Yost *et al*., 2020) and sociability (the tendency to engage in non-aggressive interactions with conspecifics; Scott *et al*., 2018). We also showed *nlg3* is required for a typical response to social isolation, but that the NLG3 protein levels are unchanged after social isolation (Yost *et al*., 2020). However, the role of *nlg3* in social recovery has not been studied.

Another molecule identified as being important for social behaviour in the fly is dopamine (DA), a monoamine neurotransmitter (Monastirioti, 1999; Waddell, 2010; Van Swinderen and Andretic, 2011). We and others have previously shown that DA modulates social space (Fernandez *et al*., 2017) and is important for a response to social isolation. Specific dopaminergic neural circuity is in part responsible for the behavioural modification after isolation (Xie *et al*., 2018) and DA levels in males are reduced after isolation (Ganguly-Fitzgerald *et al*., 2006), however the effect of DA on social recovery has not been identified. In addition, evidence suggests that *nlg3* and DA are part of a similar neurocircuitry or involved in a similar pathway responsible for behaviour regulation (Izquierdo et al., 2013; Uchigashima et al., 2016; Bariselli et al., 2018), however their dual involvement in the regulation of social space needs more attention.

In this study we examined the effect of social isolation on two measures of social behaviour, social space (Simon *et al*., 2012) and sociability (Scott *et al*., 2018). We further investigated the role of *nlg3* and DA in the recovery from social isolation. Finally, we propose a model for the integrated regulation of *nlg3* and DA in the behavioural response to the social environment.

## 2 Methods

### 2.1 Fly stocks and husbandry

All fly lines were maintained in mixed sex groups in bottles on JazzMix media or our own food made following the same recipe (brown sugar, corn meal, yeast, agar, benzoic acid, methyl paraben and propionic acid; Fisher Scientific, Whitby, ON, Canada) at 25°C, 50% relative humidity on a 12:12h light:dark cycle. All flies reared in bottles for experimental use were a maximum of 14 days old to avoid variation in behaviour resulting from older parents (Brenman-Suttner *et al*., 2018). Fly lines were obtained from the following places: CS (Canton-S) was obtained from the laboratory of Seymour Benzer in 1998, *nlg3^Def1^* are from Dr. Brian Mozer (Yost *et al*., 2020), *w;; TH-GAL4* was provided by Dr. Serge Birman, RNAi against *tyrosine hydroxylase* (w;; *UAS-THmiR-G*) was a gift from Dr. Mark Wu (Xie *et al*., 2018). All lines except w;; *UAS-ThmiR-G* were outcrossed five times to our control line, CS, to reduce variation in behaviour caused by genetic background. Crosses used to generate *TH>ThmiR-G* and their appropriate genetic controls can be found in **Supplemental Figure 1**.

### 2.2 Generation of isolation and recovery treatments

**Mated flies used in all experiments** were collected from bottles at 1 day old and remained mixed sex in new bottles for one more day (2 days total) to allow mating (see confirmation of mating status below) to avoid the effects of mating status on social space (Simon *et al*., 2012). Following, socially isolated flies were transferred to individual vials using cold anesthesia and remained singly housed for 2, 4, or 7 days. Group housed age-matched control flies were kept in mixed sex bottles for the same duration as the isolated flies, such that the group-housed control flies for isolation were 4, 6 and 9 days old respectively. To test for a recovery after social isolation, flies isolated for 7 days were transferred to mixed sex bottles for either 2 or 3 days. Group housed flies used as a control for the recovery treatment were maintained in mixed sex bottles until tested with the recovery flies, such that the group-housed control flies for isolation then recovery were 11 and 13 days old respectively. To test the effect of group housing density on social space, we sorted the group housed flies into vials mixed-sex containing 2, 6 or 16 flies and a separate uncontrolled amount (random) in a bottle.

To test for the effects of virginity on social isolation, pupae were collected from bottles and placed individually into vials and allowed to eclose over 24 hours. Isolated virgin flies remained in individual vials while group-housed virgin flies were transferred into vials containing 15 same-sex flies to maintain virginity.

**Females Mating Status at 2 and 4 days old:** one-day-old female flies were collected and maintained with males in bottles either 1 day (24 hours), or 3 days. Females were then sexed under cold anesthesia and each female was placed alone in a fresh vial. Females were scored as having been mated when 3^rd^ instar larvae were observed.

**Males Mating Status at 2 and 4 days old:** Single 1-day old male flies, sexed under cold anesthesia, were placed with one 4-day old virgin female in vials (based on the females results, more than 90% of females that age can mate). After either 1 day (24 hours), or 3 days, the males were removed, and females remained in isolation until third instar larvae were counted. For both sexes, vials with dead flies were not counted. We cannot ensure all flies have mated, even in the group housing treatment, so all flies exposed to initial group housing are used in the behaviour experiments, including the small percentage of non-mated flies.

### 2.3 Social behaviour assays

**Sociability tests** were performed at McMaster university. All other experiments took place at Western University under similar settings.

**Fly handling prior to behaviour:** Twenty-four hours prior to all behaviour assays, 15-17 flies that were group-housed (either lifelong or after a few days of recovery) were collected using cold anesthesia and placed in vials. Single housed flies were kept isolated until right before testing.

The morning of the experiment, all flies were transferred to new vials and allowed to habituate to the testing conditions of 25°C and 50% relative humidity for at least two hours. **The social space assay** was conducted under uniform light in the same room between 1:00 PM and 5:00 PM (ZT 4-8) to decrease behavioural variation linked to diel periodicity.

**The social space assay:** Performed as previously described (Simon et al., 2012;McNeil et al., 2015;Yost et al., 2020). In short: once flies have settled at their preferred inter-individual distance, a photo was taken (after 20-50 minutes, depending on their genotype). Using the open access software ImageJ (RRID:SCR_003070 - Schneider *et al*., 2012), the number of flies within 4 body lengths (∼1cm) was determined for each fly in the chamber. The number of flies within 4 body lengths was averaged using all flies in the chamber, representing one individual replicate. This metric has been used in previous studies (Xie et al., 2018; Yost et al., 2020). The routines for image analysis are publicly available (Yost *et al*., 2020). All data sets are the combination of 1-3 replicates per day and 3-5 independent days of testing. Each independent day was separated by at least a week to control for environmental factors beyond our control.

**Sociability assay:** The sociability chamber consisted of a circular arena (90 mm wide by 20 mm high) divided into 8 compartments with a hole in the center to allow flies to enter any compartment. We added to each compartment a patch of fresh food coated with a layer of grapefruit-yeast suspension (3 g yeast in 100 mL grapefruit juice) to enhance attractiveness. We modified the chamber and performance of the assay from Scott, Dworkin, and Dukas (Scott *et al*., 2018). We transferred by mouth aspiration sixteen same-sex flies to the chamber through a hole in the lid and allowed them to acclimate for one hour. Experimenters blind to treatment then counted the number of flies in each chamber and calculated an aggregation index (sample variance divided by the mean number of flies in each chamber). The variance could take values between 0 and 32. For example, the least sociable option would have 8 chambers of 2 flies each, with a variance of 0 and the corresponding aggregation index of 0. The most sociable situation would have 7 chambers of 0, and 1 chamber with all 16 flies. This would have a variance of 32, and therefore an aggregation index of 16 (32/2). We tested flies in sessions of four replicate arenas per treatment over three consecutive weeks, for twelve arenas total per treatment (total N = 96).

### 2.4 Dopamine quantification

Adults were separated by sex, and dopamine extracted from their heads using the following procedure. Extraction occurred by flash freezing flies in liquid nitrogen followed by manual decapitation and homogenization of heads in 5 mM of ammonium acetate in 90% acetonitrile using microtissue grinders (Kimble Chase, USA). The supernatant was transferred and filtered through a 0.65 µm filter (Millipore) at 4°C and samples were stored at −80 °C before LC/MS detection.

LC/MS analysis was performed using an Agilent 1260 Infinity LC system coupled to an Agilent 6230 TOF system. A XBridge C-18 column Rapid Resolution HT was used (4.6 Å∼150 mm, 3.5 μm, 600 Bar, Waters) at 25 °C and samples were eluted with a gradient of CH_3_CN (Solvent B: 90% CH_3_CN in H_2_O, containing 0.1% formic acid) in H_2_O (Solvent A: containing 0.1% formic acid). The UV lamp was set at 282 nm, the injection volume was 10 µL. The flow rate was set to 0.4 mL/min and infused into an Agilent 6230 TOF-MS through a Dual Spray ESI source with a gas temperature of 325 °C flowing at 8 L/min, and a nebulizer pressure of 35 psi. The fragmentor voltage was set to 175 V with a capillary voltage of 3500 V and a skimmer voltage of 65 V. The instrument was set in positive ESI mode, and quantification occurred using a standard curve of known dopamine concentrations (**Supplemental Figure 2**). Total ion count was extracted using Agilent MassHunter Qualitative Analysis software (version B.05.00).

### 2.5 Statistics

Data were stored in Excel files, and statistically analyzed using GraphPad Prism (RRID:SCR_002798, version 7.0a for Mac, GraphPad Software, La Jolla California USA, www.graphpad.com). We first assessed the normality and homoscedasticity of the data distribution prior to applying the appropriate parametric tests (Simple *t*-test or ANOVA or Welch-*t*-test/ANOVA when the standard deviations were different among groups). One-way and Two-way ANOVAs were used, followed by Tukey’s or Sidak’s post hoc test (as appropriate) to correct for multiple comparisons (see Table 1 for all the Two-Way ANOVA results). We used the traditional alpha level of 0.05 for all statistical tests (Wasserstein and Lazar, 2016).

**Table 1.**
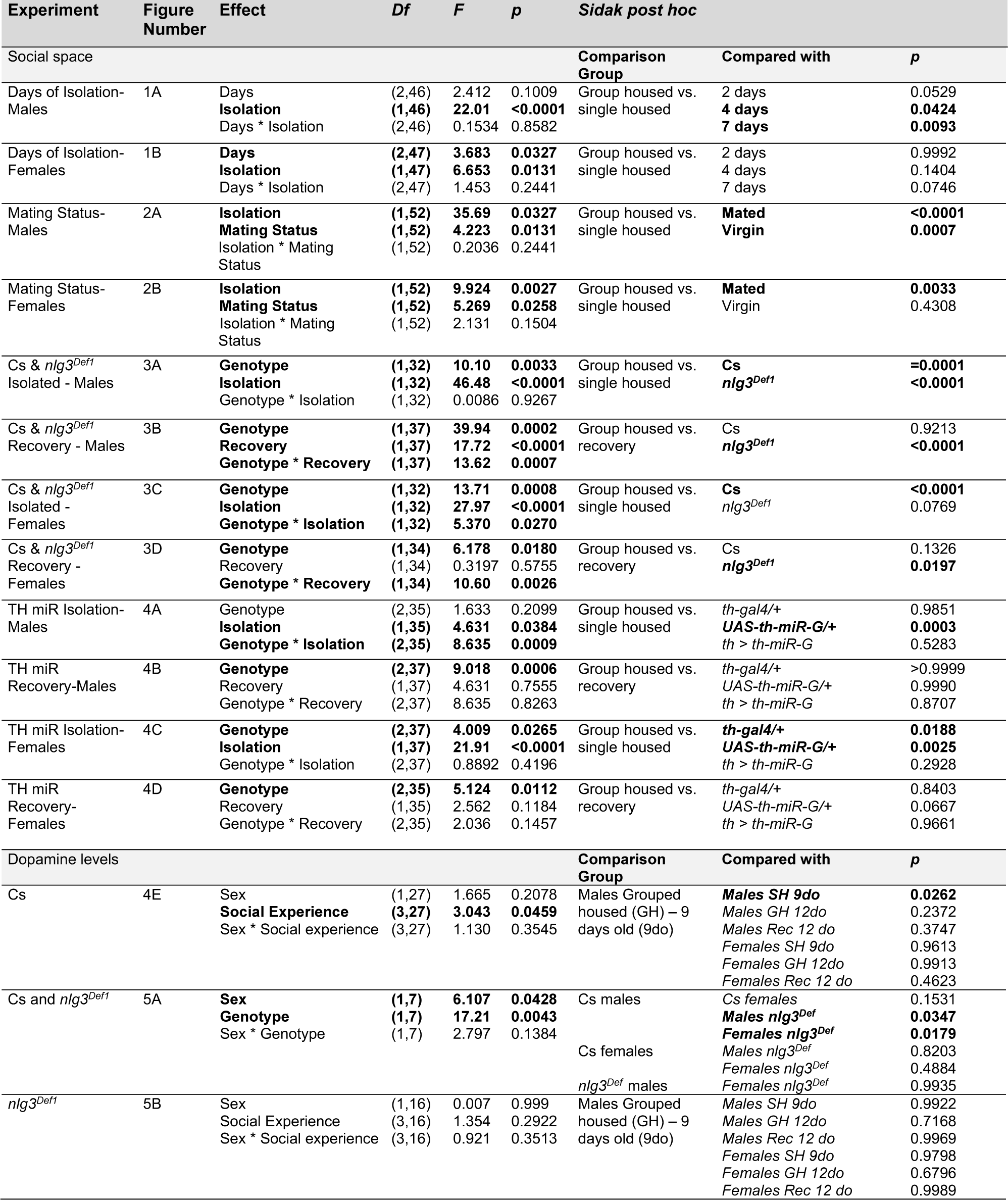
ANOVA table for all Two-way ANOVAs performed. Statistically significant results are bolded.

## 3 Results

### 3.1 Isolation for seven days leads to increased social space and decreased sociability

Seven days of social isolation leads to an increased social space (Simon *et al*. 2012). We wanted to determine whether we could reduce the duration of social isolation and still influence social space.

Because mating status influences social space, and virgin flies are further apart than mated flies (Simon *et al*., 2012), we needed to first assess how long flies should be group housed post-emergence to avoid the confounding effect of virginity on social space. We found that 101 out of 112 (90.2%) of females were mated and 55 out of 72 (76.4%) of males were able to mate after spending 4-5 days, mixed with females (**Supplemental Table 1**). In comparison, 94/103 (91.2%) of females were mated 2 days after adult emergence spent with males, and 69/81 (85.2%) of males of the same age were able to mate. The ages at which the flies were tested had no significant effect on their mating capabilities (χ²(1) = 0.022, *p*=0.88 for females, and (χ²(1) = 1.92, *p*=0.166 for males). We thus decided to expose our flies to two days of group housing post-emergence, before socially isolating them for 2, 4 or 7 days, as we now know that the same percentage of flies have mated by that age, as when they are 4-5 days. We then assessed their social space by recording the number of flies present within 4-body lengths.

Canton-S (CS, our control line) males had fewer flies within 4 body lengths after social isolation compared to group housed flies at all three time points (2, 4 or 7 days) (**Figure 1A**; Two-way ANOVA results in **Table 1**; Effect of isolation: *F_1,46_* = 22.01, *p* < 0.0001). There was no effect of 2 days of social isolation in CS females. Only after 4 and 7 days of isolation did CS females have fewer flies within 4 body lengths and the increase in social space was larger with increasing number of days flies were isolated (**Figure 1B**; Two-way ANOVA results in **Table 1**; Effect of isolation: *F_1,47_* = 6.653, *p* = 0.0131; Effect of days of isolation: *F_2,47_* = 3.683, *p* = 0.0327). The effect of isolation after 7 days was similar to the effects found in earlier studies (Simon *et al*., 2012). Moving forward we used seven days of isolation for the rest the experiments as seven days had the largest increase in social space in males and females.

**Figure 1.**
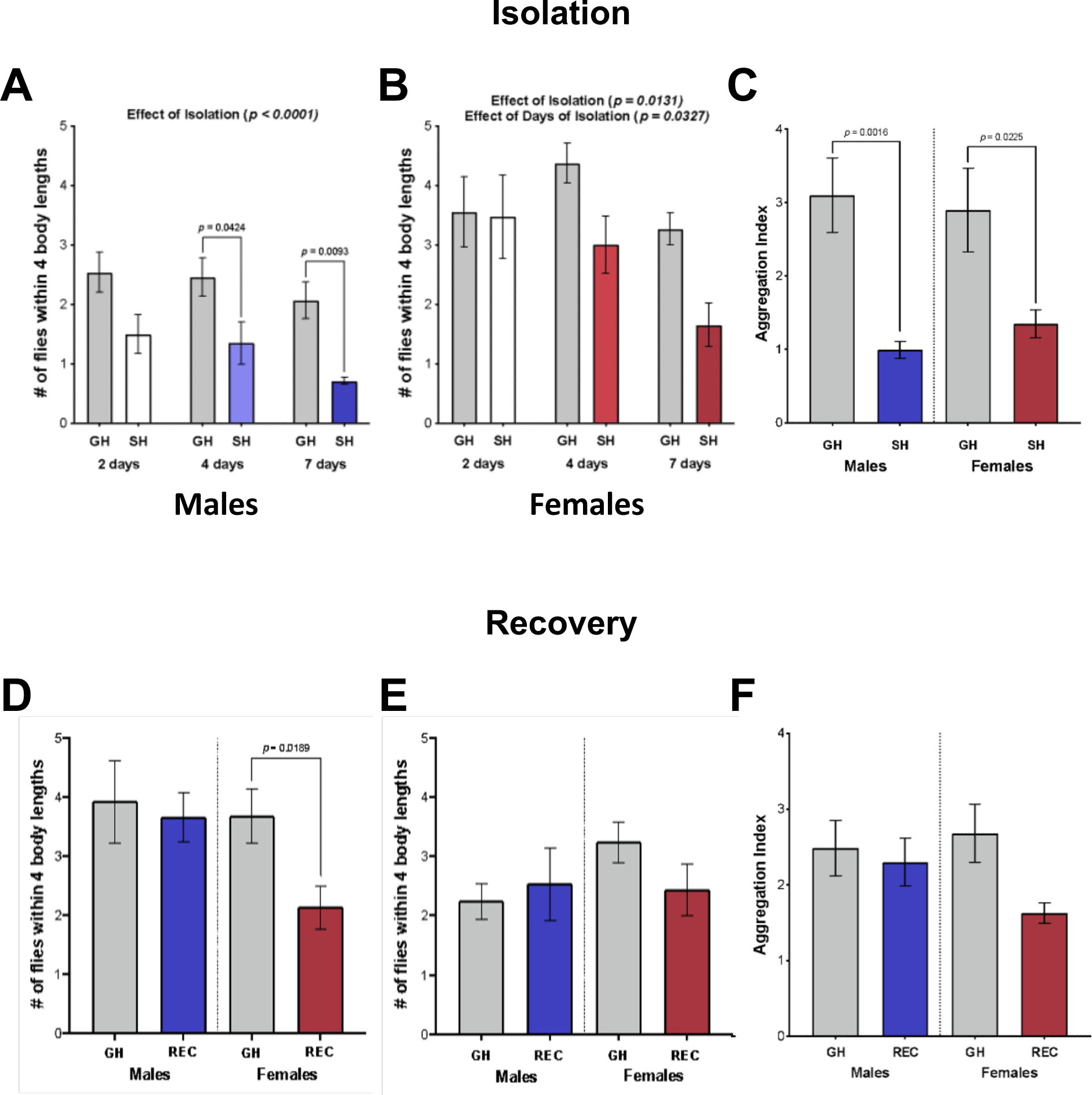
Isolation for seven days leads to increased social space and decreased sociability, but flies recover after 3 days of group housing. **A-B:** Social space presented in terms of mean number of flies within 4 body lengths +/− s.e.m. in males (A) and females (B) after 2, 4 or 7 days of isolation. **A.** Male flies had fewer flies within 4 body lengths after 2, 4 and 7 days of isolation (Two-way ANOVA and Sidak post hoc test - *F*_1,46_ = 22.01, *p* < 0.0001). **B.** Females had fewer flies within 4 body lengths after isolation and that number decreased further with increasing days of isolation (Two-way ANOVA - Effect of Isolation: *F*_1,47_ = 6.653, *p* = 0.0131; Effect of Days of Isolation: *F*_2,47_ = 3.687, *p* = 0.0367). **C**. Aggregation index in males and females after 7 days of isolation. Both males (Welch’s T-test: t_12.08_ = 4.039, *p* = 0.0016) and females (Welch’s T-test: *t*_13.37_ = 2.578, *p* = 0.0225) had a lower aggregation index after 7 days of isolation. **D-E:** Males and females after 2 (D) and 3 (E) days of group housing (recovery) following 7 days of isolation. **D.** Males showed no difference in the number of flies within 4 body lengths after 2 days of group housing compared to males continuously group housed. However, females had fewer flies within 4 body lengths compared to females group housed continuously (Welch’s T-test: *t*_15.13_ = 2.626, *p* = 0.0189). **E.** Both males and females were not different in the number of flies within 4 body lengths comparing group housed and recovery flies (One-way ANOVA: *F*_3,28_ = 0.9434, *p* = 0.4329). **F.** Sociability presented as the mean aggregation index +/− s.e.m. of males and females group housed and recovery. Males and females did not differ in aggregation index between group housed and recovery flies (One-way ANOVA: *F*_3,44_ = 2.102, *p* = 0.1136). N = 7-12 for all treatments. \**p* < 0.05. \*\*\*\**p* < 0.0001. Grey shade: Group Housed (GH); blue shades: single housed (SH) and recovery from social isolation (REC) males; red shades: SH and REC females.

We next used another measure of social behaviour, sociability (Scott *et al*., 2018), to examine the effect of isolation. Both males and females had a lower aggregation index in isolated flies compared to those group housed (**Figure 1C, Table 2**; Males – Welch’s T-test: *t*_12.08_ = 4.039, *p* = 0.0016; Females – Welch’s T-test: *t*_13.37_ = 2.578, *p* = 0.0225).

**Table 2.** All t-tests performed. Statistically significant results are bolded

| Experiment | Figure Number | Comparison | Df | t | p |
| --- | --- | --- | --- | --- | --- |
| Sociability Isolation – Males | 1C | Group housed vs. single housed males | **12.08** | **4.039** | **0.0016** |
| Sociability Isolation – Females | 1C | Group housed vs. single housed females | **13.37** | **2.578** | **0.025** |
| Social Space Recovery 2 day – Males | 1D | Group housed vs. recovery males | 13.11 | 0.3097 | 0.7617 |
| Social Space Recovery 2 day – Females | 1D | Group housed vs. recovery females | **15.13** | **2.2626** | **0.0189** |

### 3.2 Males and females recover from isolation after three days of group-housing

As recovery from isolation has been noted in various organisms including *Drosophila melanogaster* (see introduction), we tested whether social space and sociability could also be recovered in flies that were group housed following isolation. We started with 2 days of group housing following isolation and found that males were not different in flies within 4 body lengths indicating they had recovered in social space, however females still had a lower number of flies within 4 body lengths in the recovery compared to group housed flies (**Figure 1D, Table 2**; Welch’s T-test: *t*_15.13_ = 2.626, *p* = 0.0189). Because females had not fully recovered after two days of group housing post isolation, we tested social space in flies that had three days of group housing after seven days of isolation. Again, males were not different in flies within 4 body lengths, however females were now also not different in flies within 4 body lengths in recovery and group housed flies indicating that females also can recover from isolation but take longer than males (one more day in this case – **Figure 1E, Table 3**). A period of three days of group housing following seven days of isolation was then used for all other experiments investigating recovery. Finally, we tested sociability in males and females after 3 days of recovery. Males and females did not differ in aggregation index in recovery and group housed flies, indicating that both sexes recovered (**Figure 1F, Table 3**).

**Table 3.** ANOVA table for all One-way ANOVAs performed. Statistically significant results are bolded.

| Experiment | Figure Number | Df | F | p | Sidak post hoc |  |  |
| --- | --- | --- | --- | --- | --- | --- | --- |
|  |  |  |  |  | Comparison Group | Compared with | p |
| Social Space Recovery – Males & Females | 1E | (3,28) | 0.9434 | 0.4329 | Group housed Males | Recovery Males | 0.8702 |
|  |  |  |  |  | Group housed Females | Recovery Females | 0.3881 |
| Sociability Recovery – Males & Females | 1F | (3,44) | 2.102 | 0.1136 | Group housed Males | Recovery Males | 0.8841 |
|  |  |  |  |  | Group housed Females | Recovery Females | 0.2599 |
| Group Housed Density – Males | 2C | (4,40) | 30.87 | <0.0001 | Single Housed | 2 | <0.0001 |
|  |  |  |  |  |  | 6 | <0.0001 |
|  |  |  |  |  |  | 16 | <0.0001 |
|  |  |  |  |  |  | Random | <0.0001 |
| Group Housed Density – Females | 2C | (4,40) | 31.35 | <0.0001 | Single Housed | 2 | <0.0001 |
|  |  |  |  |  |  | 6 | <0.0001 |
|  |  |  |  |  |  | 16 | <0.0001 |
|  |  |  |  |  |  | Random | <0.0001 |

### 3.3 Mating status does not alter social space in response to isolation

We have previously reported that virgin flies have an increased social space (Simon *et al*., 2012) as do isolated flies (Yost *et al*., 2020 and Figure 1), however we have not tested virgin flies in response to isolation, so we next tested the effect of isolation on social space in mated and virgin flies. There were less flies within 4 body lengths after isolation in both mated and virgin males (**Figure 2A**; Two-way ANOVA results in **Table 1**; Effect of isolation: *F_1,52_ = 34.69, p < 0.0001*). In addition, virgin males had less flies within 4 body lengths when group and single housed compared to mated group and single housed (**Figure 2A**; Two-way ANOVA results in **Table 1**; Effect of mating status: *F_1,52_ = 4.223, p = 0.0449*). In females we found that in mated flies, single housed flies had less flies within 4 body lengths than group housed individuals (**Figure 2B**; Two-way ANOVA results in **Table 1**; Effect of isolation: *F_1,52_ = 9.924, p = 0.0027*), however, group and single housed virgins had similar flies within 4 body lengths (*p* = 0.4308). Similar to males, virgin females had less flies within 4 body lengths when group and single housed compared to mated group and single housed **Figure 2B**; Two-way ANOVA results in **Table 1**; Effect of mating status: *F_1,52_ = 5.269, p = 0.0258*).

**Figure 2.**
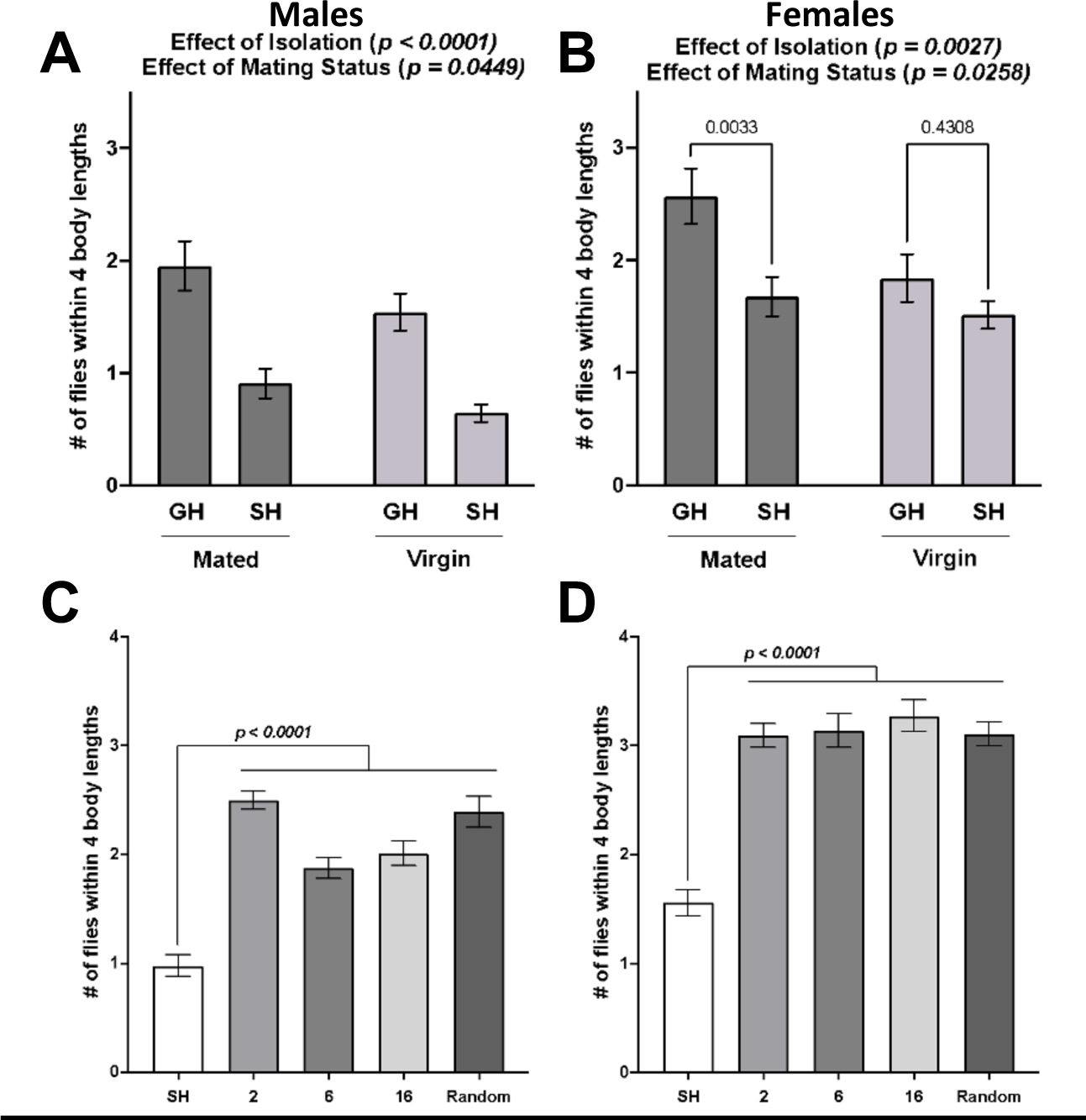
Mating status does not change social space in response to isolation and group housed with a single fly is enough to avoid a larger social space. **A-B:** Number of flies within 4 body lengths in males (A) and females (B) group and single housed while mated or virgin. **A.** Male flies had less flies within 4 body lengths after isolation (Two-way ANOVA – Effect of isolation: *F*_1,52_ = 34.69, *p* < 0.0001) regardless of mating status. Males had less flies within 4 body lengths when virgin compared to mated flies (Two-way ANOVA – Effect of mating status: *F*_1,52_ = 4.223, *p* = 0.0449). **B.** Female flies had less flies within 4 body lengths after isolation (Two-way ANOVA – Effect of isolation: *F*_1,52_ = 9.924, *p* = 0.0027) regardless of mating status. Females had less flies within 4 body lengths when virgin compared to mated flies (Two-way ANOVA – Effect of mating status: F_1,52_ = 5.269, p = 0.0258). **C-D:** Number of flies within 4 body lengths in males (C) and females (D) reared single housed or group housed with 2, 6, 16 or a random number of flies. Single housed flies had less flies within 4 body lengths than group housed flies that were reared with any number of individuals in males (C; One-way ANOVA with Holm-Sidak post hoc: *F*_*4*,40_ = 30.87, *p* < 0.0001) and females (D; One-way ANOVA with Holm-Sidak post hoc: *F*_*4*,40_ = 31.35, *p* < 0.0001). N = 9-13 for all treatments. GH: Group housed, SH: Single housed. Bars: mean +/− s.e.m.

### 3.4 Group housed density does not alter social space

We next tested what happens when group housed flies are reared at different densities ranging from 2 flies up to an uncontrolled or random number of flies. We found that single housed flies have less flies within 4 body lengths compared to group housed flies at all densities in males (**Figure 2C, Table 3**; One-way ANOVA with Holm-Sidak *post hoc*: *F*_*4*,40_ = 30.87, *p* < 0.0001) and females (**Figure 2D, Table 3**; One-way ANOVA with Holm-Sidak *post hoc*: *F*_*4*,40_ = 31.35, *p* < 0.0001).

### 3.5 The lack of *nlg3* does not prevent from recovering from social isolation

We have previously shown that *nlg3* is important for a typical social space response to isolation (Yost *et al*., 2020). We wondered whether *nlg3* is also important for the recovery from social isolation. In males, both CS and *nlg3^Def1^* had less flies within 4 body lengths in single housed compared to group housed flies and *nlg3^Def1^* had less flies within 4 body lengths than CS (**Figure 3A**, Two-way ANOVA results in **Table 1**; Effect of social experience: *F_1,32_ = 46.48, p < 0.0001;* Effect of genotype: *F_1,32_ = 10.10, p = 0.0033*). When testing the recovery, CS males did not differ in the number of flies within 4 body lengths, however *nlg3^Def1^* males had increased flies within 4 body lengths in the recovery treatment compared to group housed *nlg3^Def1^*flies (**Figure 3B**, Two-way ANOVA results in **Table 1**; Effect of social experience: *F_1,37_ = 17.72, p = 0.0002;* Effect of genotype: *F_1,37_ = 34.94, p > 0.0001*; Interaction of social experience and genotype: *F_1,37_ = 13.62, p = 0.0007*).

**Figure 3:**
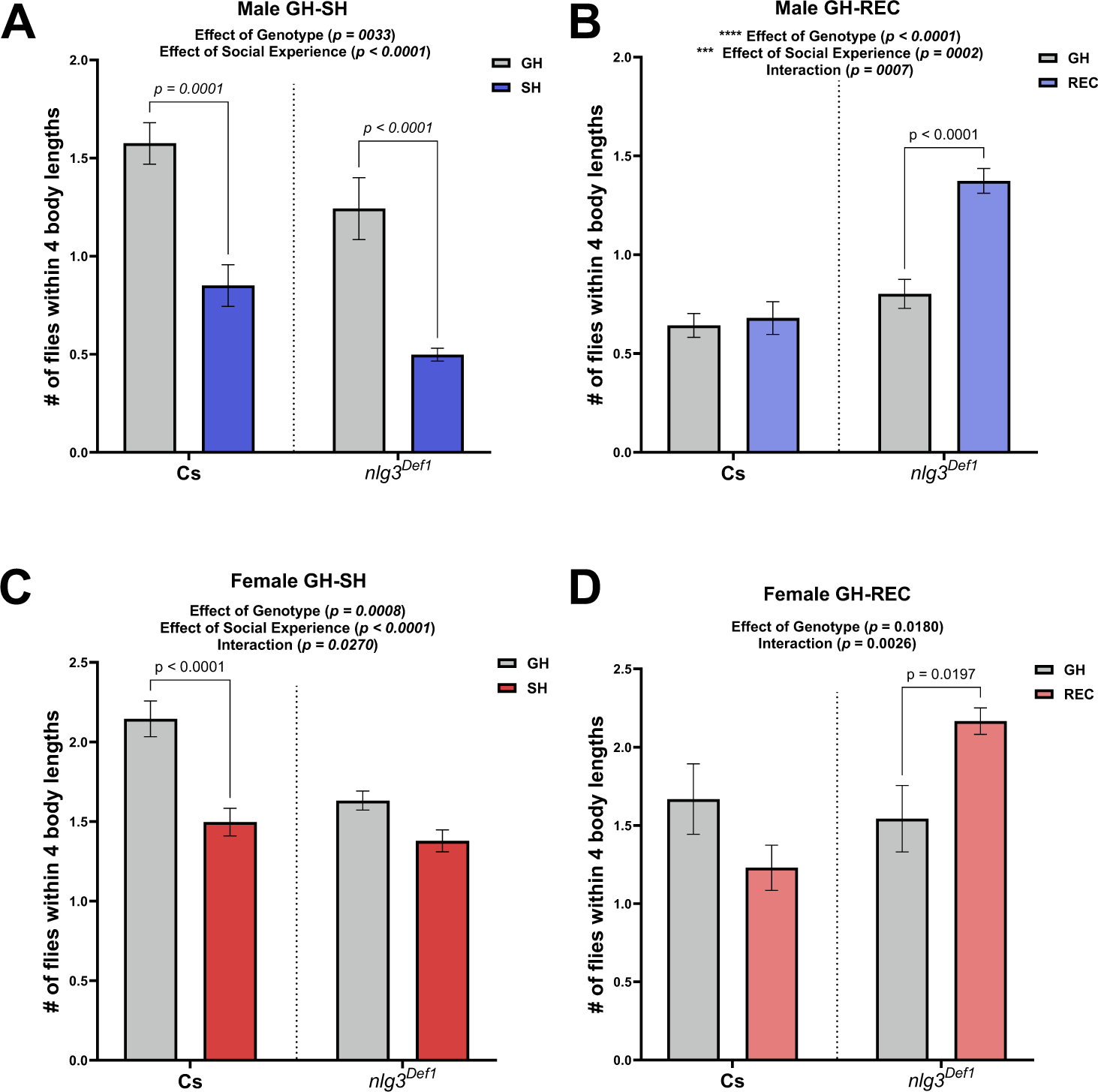
*nlg3^Def1^* flies have decreased social space after a recovery from isolation. **A-B:** Number of flies within 4 body lengths in males group and single housed (A) and group housed with a recovery (B). **A.** Single housed male flies had decreased flies within 4 body lengths compared to group housed flies in both CS and *nlg3^Def1^* (Two-way ANOVA - Effect of social experience: *F*_1,32_ = 46.48, *p* < 0.0001) and *nlg3^Def1^*flies had less flies within 4 body lengths in both group housed and single housed (Two-way ANOVA - Effect of genotype: *F*_1,32_ = 10.10, *p* = 0.0033). **B.** The number of flies within 4 body lengths for CS males was not different in group housed vs recovery flies, however *nlg3^Def1^*flies had an increased number of flies within 4 body lengths in recovery compared to group housed flies (Two-way ANOVA - Effect of social experience: *F*_1,37_ = 17.72, *p* = 0.0002; Effect of genotype: *F*_1,37_ = 34.94, *p* < 0.0001; Interaction of social experience and genotype: *F_1,37_* = 13.62*, p* = 0.0007). **C-D:** Number of flies within 4 body lengths in group and single housed females (C) and group housed with a recovery females (D). **C.** CS females had less flies within 4 body lengths in single housed compared to group housed flies, however *nlg3^Def1^* flies were not different between group and single housed flies (Two-way ANOVA - Effect of social experience: *F*_1,32_ = 27.97, *p* < 0.0001; Effect of genotype: *F*_1,32_ = 13.71, *p* = 0.0008; Interaction of social experience and genotype: *F_1,32_* = 5.370*, p* = 0.0270). **D.** The number of flies within 4 body lengths for CS females was not different in group housed vs recovery flies, however *nlg3^Def1^*females had an increased number of flies within 4 body lengths in recovery compared to group housed flies (Two-way ANOVA - Effect of social experience: *F*_1,34_ = 0.3197, *p* = 0.5755; Effect of genotype: *F*_1,34_ = 6.178, *p* = 0.0180; Interaction of social experience and genotype: *F_1,34_* = 10.60*, p* = 0.0026). N = 7-12 for all treatments. GH: Group housed. GH: Group housed, SH: Single housed, REC: Recovery.

In females, CS had less flies within 4 body lengths when single housed, however *nlg3^Def1^* females had no difference in the number of flies within 4 body lengths compared to group housed flies (**Figure 3C**, Two-way ANOVA results in **Table 1**; Effect of social experience: *F_1,32_ = 27.97, p > 0.0001;* Effect of genotype: *F_1,32_ = 13.71, p = 0.0008*; Interaction of social experience and genotype: *F_1,32_ = 5.370, p = 0.0270*). In the recovery from isolation, CS flies were not different in the number of flies within 4 body lengths compared to flies always group housed however *nlg3^Def1^* flies had increased number of flies within 4 body lengths in the recovery treatment compared to group housed flies (**Figure 3D**, Two-way ANOVA and Tukey *posthoc* results in **Table 1**; Effect of social experience: *F_1,34_ = 0.3197, p = 0.5755;* Effect of genotype: *F_1,34_ = 6.178, p = 0.0180*; Interaction of social experience and genotype: *F_1,34_ = 10.60, p = 0.0026*).

This recovery experiment is repeated in **Supplemental Figure 3** (those data were in fact collected at the same time as the isolation treatments published in Yost *et al*., 2020).

Combined, the data indicate that *nlg3* is not required for recovery to occur.

### 3.6 Dopamine is required for a response to isolation and decreases after isolation in a sex-specific manner

Dopamine is important not only for social space in both sex (Fernandez *et al*., 2017) but also for the response to social isolation in males (Xie *et al*., 2018). We tested to see if we could recapitulate these results after isolation and determine if dopamine was important for the recovery from isolation. We drove an RNAi against the gene encoding for Tyrosine Hydroxylase (*TH*), the rate limiting enzyme for dopamine biosynthesis (*TH*-RNAi, specifically the *UAS*-*THmiR-G*) using a *TH-gal4* driver.

In males *TH>THmiR-G*, there was no effect of social experience. We did observe a significant reduction in the number of flies within 4 body lengths with isolation in flies with *TH-*Gal4/+ and *UAS-THmiR-G*/+ (**Figure 4A**; Two-way ANOVA results in **Table 1**; Effect of isolation: *F_1,42_* = 19.13, *p* < 0.0001; Genotype and isolation interaction: *F_2,35_* = 8.635, *p* = 0.0009). When investigating the recovery, we observed no difference in any genotype between group housed and recovery treatments (**Figure 4B**; Two-way ANOVA results in **Table 1**).

**Figure 4.**
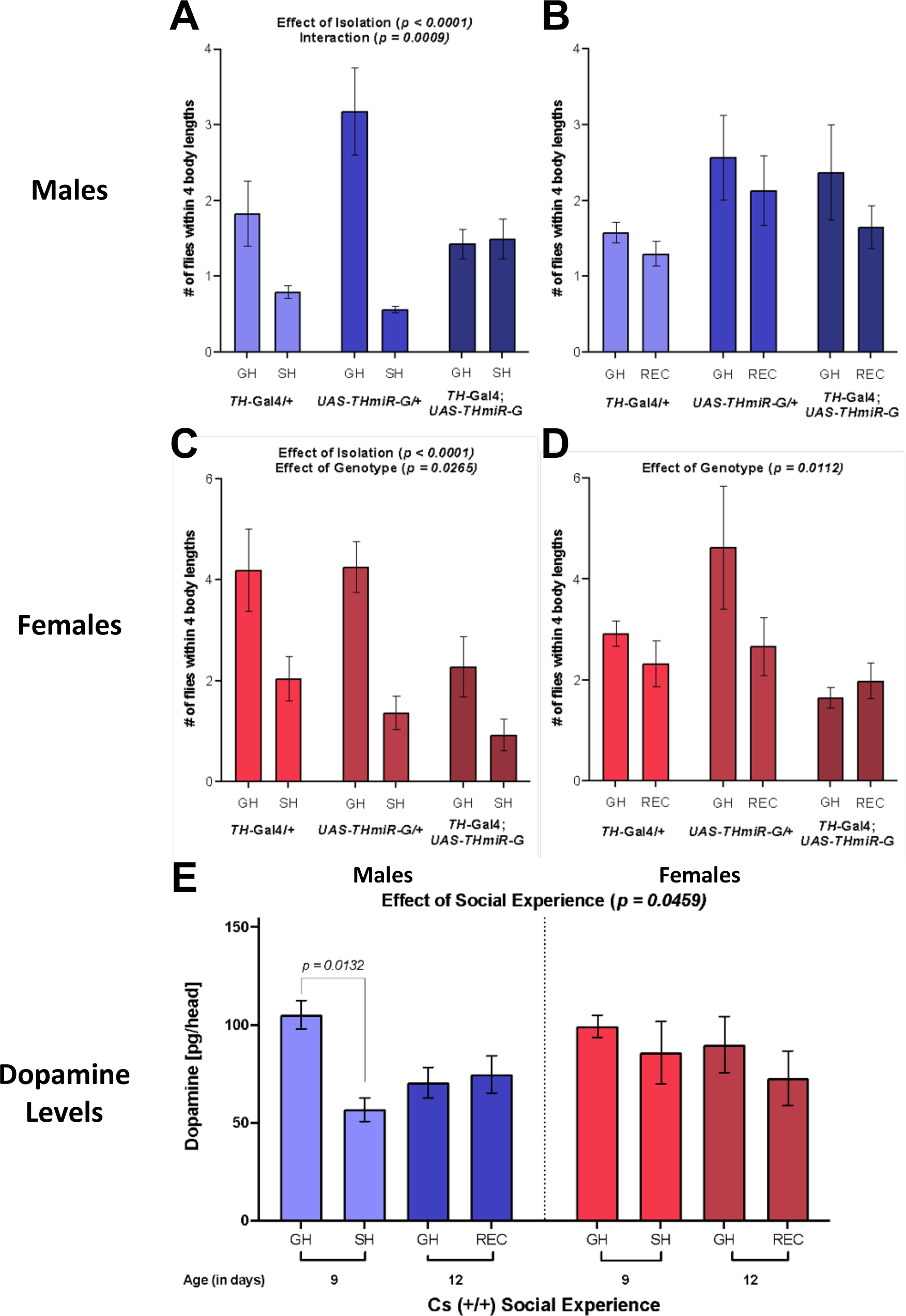
Dopamine is important for a response to social isolation and decreases after isolation in males but not females. Social space measured in terms of number of flies within 4 body lengths. Bars: mean +/− s.e.m. **A-B:** Social space in isolated (A) and recovered (B) male flies. **A.** The social space in male *TH > THmiR-G* was not different in isolated vs group housed flies, however *TH-GAL4/+* and *UAS-THmiR-G/+* males had lower flies within 4 body lengths when isolation (Two-way ANOVA - Effect of Isolation: *F*_1,43_ = 19.13, *p* < 0.0001; Interaction between isolation and genotype: *F*_2,42_ = 8.254, *p* = 0.0009). **B.** Male recovery flies in all genotypes were not different compared to group housed males. **C-D:** Social space in isolated (C) and recovered (D) female flies. **C.** All three genotypes had a decrease in the number of flies within 4 body lengths after isolation and *TH > THmiR-G* flies had the lowest number of flies within 4 body lengths compared to their genetic controls (Two-way ANOVA – Effect of Isolation: *F*_1,37_ = 21.91, *p* < 0.0001; Effect of Genotype: *F*_*2*,37_ = 4.009, *p* = 0.0265). **D.** All three genotypes were similar when comparing group housed vs. recovery treatments, however *TH > THmiR-G* females had a less flies within 4 body lengths compared to the controls (Two-way ANOVA – Effect of Genotype: *F*_2,35_ = 5.124, *p* = 0.0112). **E.** Dopamine levels (pg/head) in males decreased in isolated compared to group housed flies but do not differ in recovery flies compared to group housed in males or females (Two-Way ANOVA – Effect of Social Experience: *F*_3,27_ = 3.043, *p* = 0.0459, but the only significant difference after correcting for multiple comparison through a Dunnett test is indicated on graph, *p*=0.0132). Appropriate age comparisons were made (see methods for details). \**p* < 0.05. **** *p* < 0.0001. Bars: mean +/− s.e.m. N=5-9 for all treatments in social space. N=4 for dopamine quantification. GH: Group Housed, light grey shade; SH: Single Housed, dark grey shade; REC: Recovery from social isolation, medium grey shade.

In females, we observed a decrease in the number of flies within 4 body lengths for all three genotypes after isolation and a decrease in *TH>THmiR-G* flies compared to their genetic controls (**Figure 4C**; Two-way ANOVA results in **Table 1**; Effect of genotype: *F_2,37_* = 4.009, *p* = 0.0265; Effect of isolation: *F_1,37_* = 21.91, *p* < 0.0001). Female *TH>THmiR-G* also displayed a decreased number of flies within 4 body length; however, no genotype was different comparing group housed to recovery treatments (**Figure 4D**; Two-way ANOVA results in **Table 1**; Effect of genotype: *F_2,35_* = 5.124, *p* = 0.0112).

To establish the importance of dopamine in a response to isolation and recovery, we performed LC/MS on the heads of males and females to quantify changes in dopamine levels after isolation and recovery. After isolation, males had decreased dopamine levels while females’ dopamine levels remained similar to those of group housed flies. Further, male dopamine levels returned to a similar level as group housed after a recovery period. Females’ dopamine levels remained unchanged (**Figure 4E**; Two-way ANOVA results in **Table 1**; Effect of social experience: *F_3,27_* = 3.043, *p* = 0.0459).

Taken together, these results indicate that dopamine is important for social space in response to isolation and recovery in males but not females, and that dopamine levels are influenced by previous social experience in a sex-specific manner.

### 3.7 *nlg3^Def1^* flies have less dopamine, but its levels are unchanged after isolation

To determine if an interaction was occurring between DA and *nlg3* we tested DA levels in group housed CS and *nlg3^Def1^.* In both males and females, DA was decreased in *nlg3^Def1^* compared to CS and females had lower DA levels compared to males (**Figure 5A**; Two-way ANOVA results in **Table 1**; Effect of sex: *F_1,7_* = 6.107, *p* = 0.0428; Effect of genotype: *F_1,7_* = 17.21, *p* = 0.0043). However, when we tested the effect of social experience, no difference in DA levels was observed between *nlg3^Def1^* group housed, isolated and recovery males and females (**Figure 5B**; Two-way ANOVA results in **Table 1**).

**Figure 5.**
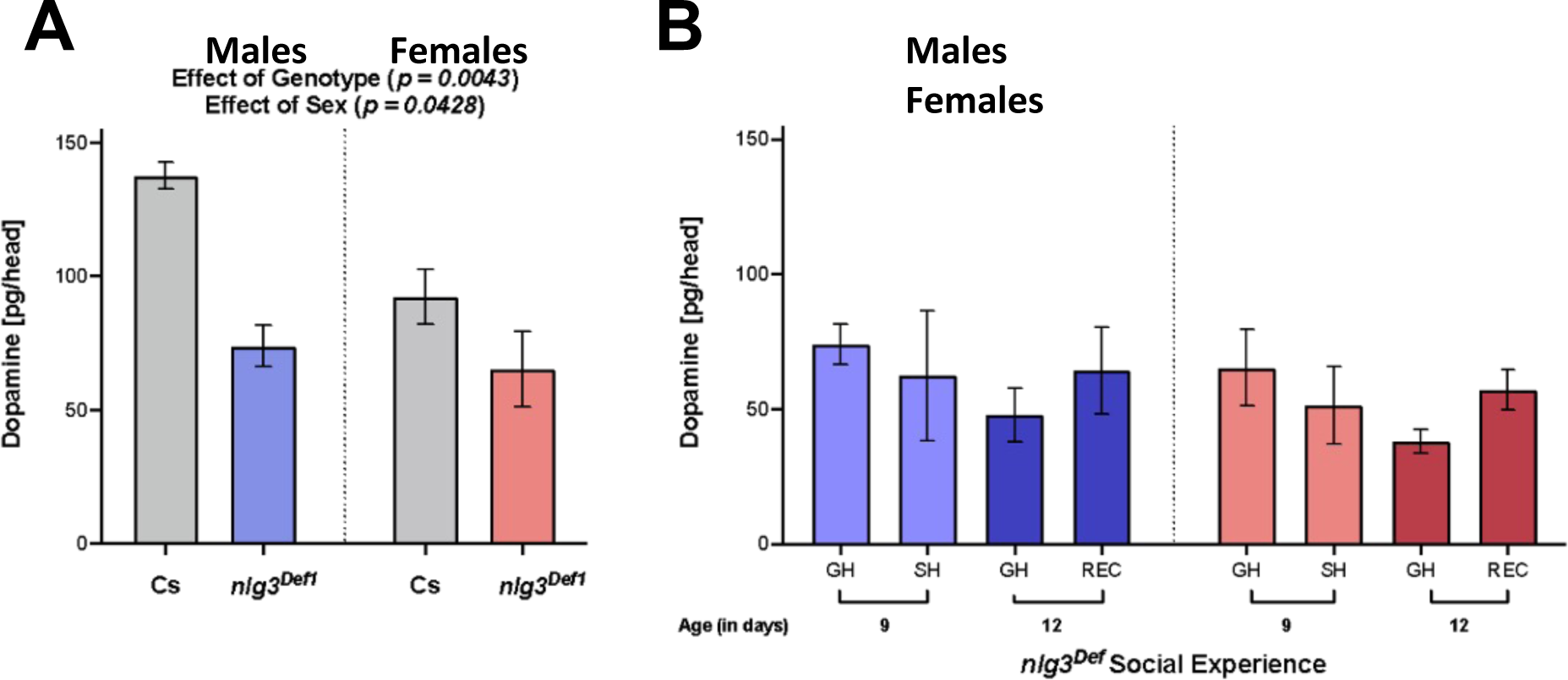
*nlg3^Def1^*flies have less dopamine, and those levels are not altered by social experience. **A.** DA levels (pg/head) in CS (grey shade) and *nlg3^Def1^*males (blue shade) and females (red shade). *nlg3^Def1^* flies had less DA when group housed and isolated compared to group housed CS, with females showing a lesser amount overall (Two-way ANOVA, Effect of Genotype: *F*_1,7_= 17.21, *p* = 0.0043 and Effect of Sex: *F*_1,7_ = 6.107, *p* = 0.0428). **B.** DA levels (pg/head) in male (blue shade) and female (red shade) group housed and recovery *nlg3^Def1^* flies are unchanged with social experience. Appropriate age comparisons were made (see methods for details). Bars: mean +/− s.e.m. N=3-6 for all treatments. GH: Group Housed. SH: Single Housed. REC: Recovery from social isolation. ***P < 0.001. **P < 0.01. *P < 0.05.

## 4 Discussion

Here we report the effects of social isolation on social space and sociability. In addition, for the first time, we report the effect of social recovery on the fly social behaviour. Social space was increased, and sociability decreased after isolation but was recovered after three days of group housing following isolation. We also show that mating status affected the response to isolation in a sex-specific and that as soon as 2 flies or more are present, the size of the group prior to assaying social space as no effect on social spacing. In addition, we show that an autism candidate gene, *nlg3,* important for the response to social experience, is not required for recovery from social isolation to occur. We also further confirm the role of dopamine in social space. But we show for the first time the sex-specific role of DA in social spacing: DA is not only required for a response to isolation but also for recovery from isolation in males, but not females. Furthermore, DA levels also respond to the social environment by decreasing in males after isolation and returning to normal levels when recovery occurs, but not in females. Finally, we show that a *nlg3* loss of function mutant, *nlg3^Def1^*, inhibits a change in DA in males and females, with no further changes added with social isolation.

We first looked at how long flies needed to be isolated to effect social space. We saw a sex-specific response to isolation where males only needed two days of isolation while females needed closer to seven days. The females were less effected in the early days of isolation and seemed to be more resilient to the effects of isolation than males (**Figure 1A & B**). Our results for both males and females isolated for seven days phenocopy the results of Simon *et al*., (2012). With regard to recovery, males also responded faster, and recovered in two days versus three days in females (**Figure 1D & E**). In summary, although males were affected sooner by isolation, they were able to recover faster than females. Finally, we showed that sociability is reduced in males and females but recovers after group housing (**Figure 1C & F**), indicating that isolation is affecting multiple social behaviours in the fly.

As we have previously shown that mating status and isolation are important for social space (Simon et al., 2012; Yost et al., 2020), we investigated the role of mating status in social space in response to isolation. In both males and females, social space was increased in virgin group housed flies compared to mated group housed flies as previously reported (Simon et al., 2012; **Figure 2A**). Both virgin and mated male flies had increased social space after isolation. We then wanted to know if the number of flies present while group housed affected social space. We tested single housed flies and flies reared group housed with 2, 6, 16, and an uncontrolled random number of flies. The isolated flies had increased social space as expected (**Figure 2C & D**). Group housed flies had lower social space than isolated flies, however, the social space was similar amongst all the densities indicating the presence of even one other fly is enough to avoid the negative consequences of isolation on social space.

Next, we looked at social space in the Drosophila homolog of an autism candidate gene, *nlg3*, after isolation and recovery. We showed that males CS and *nlg3^Def1^* flies had increased social space after isolation. We have previously reported a diminished response to isolation in *nlg3^Def1^* flies (Yost *et al*., 2020), however this time, we observed that male responded to isolation similarly to their genetic controls (**Figure 3A**). However, isolated *nlg3^Def1^* females did not, similarly to what we had reported previously (**Figure 3C**; Yost *et al*., 2020). When testing the recovery after isolation, male and female CS flies had similar social space to flies always group housed, indicating recovery had occurred (**Figure 3B, D**). Interestingly, in *nlg3^Def1^* flies, both male and females had decreased social space in the recovery treatment, which appears as an over compensatory response. In an independent repeat of those data, both CS and *nlg3^Def1^* flies recovered similarly from isolation (**Supplemental Figure 3**). In both cases, lack of *nlg3* did not prevent the recovery from social isolation.

Using an RNAi against tyrosine hydroxylase, we were able to show that DA is important for a response to isolation in males since the isolated flies did not respond to isolation and even had a slightly closer social space in isolated compared to group housed flies (**Figure 4A**). A similar result was reported by Xie *et al*. (2018), where group housed flies acted as if isolated and isolated flies acted as if group housed. We do not see as strong of effect as those authors reported, but we also used a pan-TH driver, while Xie *et al*. (2018) focused on a few TH cells only. Our work and that of Xie *et al*. (2018) demonstrate the importance of DA in males in the response to the social environment. However, we show for the first time that females do not require DA for a response to isolation and recovery to occur (**Figure 4C & D**). Another neurotransmitter or neuromodulator could be at play in females, in response to social experience. It has been previously shown that both males and females respond similarly to reduced DA levels, when group housed, with an increase in social space (Fernandez *et al*., 2017). But DA might not be the neurotransmitter involved in responding to social experience in females. When we isolated our control line, CS, we confirmed that there is a decrease in DA in males but not females and that the DA levels in males returned to the level of group housed flies after recovery (**Figure 4E**). Ganguly-Fitzgerald *et al*. (2006) also saw a decrease in DA in males after isolation. Taken together these results indicate DA in males is extremely important for social space in response to isolation, but not in females. Finally, we report that DA levels in *nlg3^Def1^* remain unchanged after isolation, providing further evidence that *nlg3* and DA are both contributing to the modulation of social space after isolation (**Figure 5A & B**). In both CS and *nlg3^Def1^,* decreases in DA were observed between 9- and 12-day old flies regardless of social experience, consistent with age related decreases in DA previously reported (Neckameyer et al., 2000).

As for recovering from social isolation, what does this mean for dopamine and an autism related gene? Dopamine in males is required for a response to the environment, however the flies need *nlg3* for DA levels to change in response to the social environment and subsequently modulate behaviour. Furthermore, DA levels decrease in response to social isolation in males, but the NLG3 protein itself is not responding to social experience (Yost *et al*., 2020). We predict that *nlg3* and DA are part of a pathway responsible for the modulation of social behaviour after isolation in males. *Nlg3* would be required downstream of DA, for proper DA signaling in response to the social environment, with some feedback regulation, since DA did not change after isolation when *nlg3* was absent. Females do require *nlg3* for a typical response to isolation (Yost *et al*., 2020), but other neurotransmitters or neuromodulators, responsive to social experience in females, may be interacting with *nlg3* (**Figure 6**). A role of Nlgs in DA signaling, and their common influence on behaviour, including social behaviour, has been observed in mice (Rothwell *et al*., 2014, Bariselli *et al.,* 2018), the northern swordtail fish (Wong and Cummings, 2014) and the worm *C. elegans* (Izquierdo *et al*., 2013). What remains unknown is whether Nlg3 is directly involved in the post-synaptic recruitment of one of the dopamine receptors in flies, as has been suggested, based on work in mice, by Uchigashima *et al*. (2016). Finally, whether *Cyp20* is also involved in the response to social experience, in both males and females, to modulate changes in social space and sociability still needs to be investigated (Wang *et al.,* 2008).

**Figure 6:**
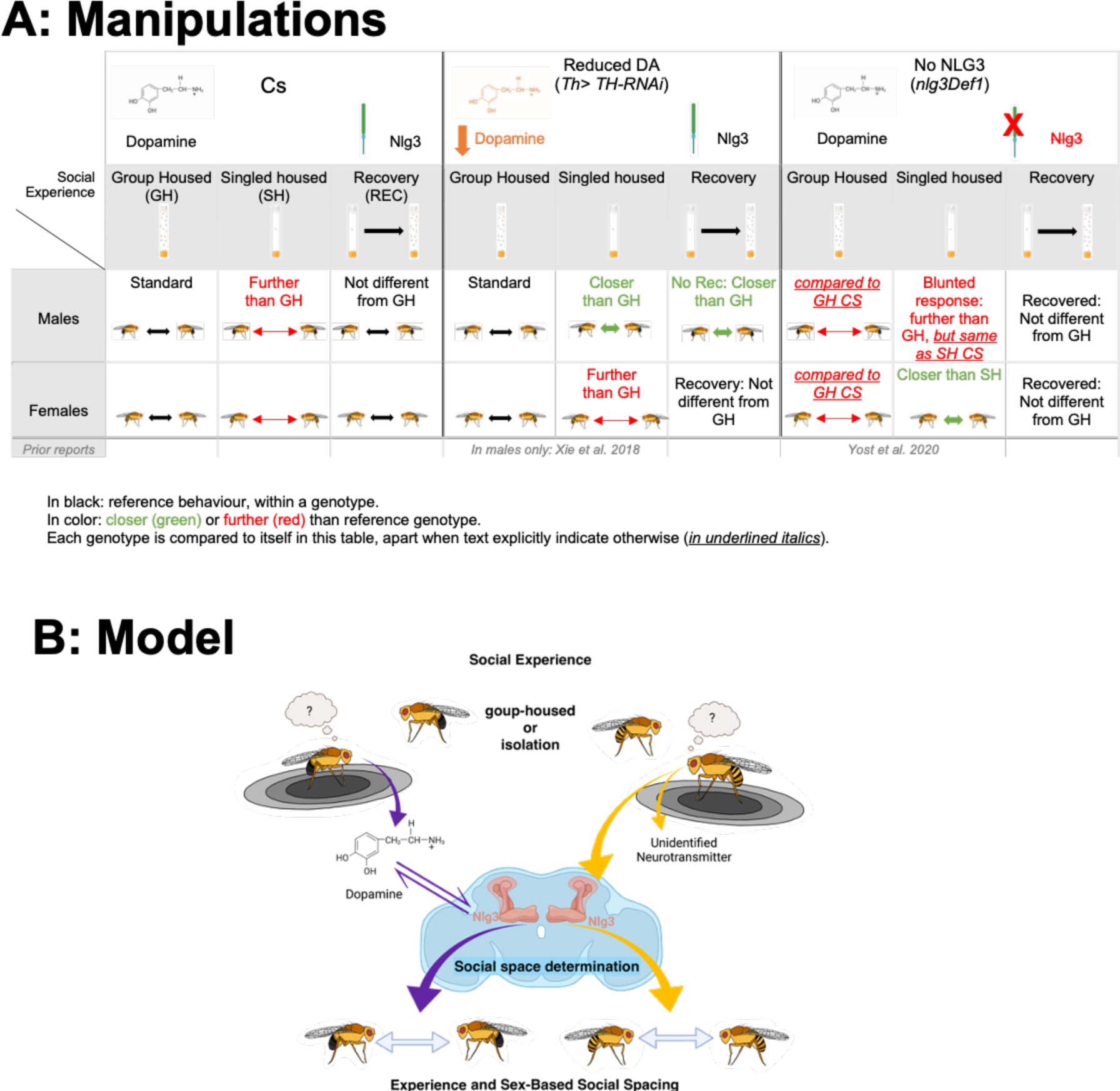
A. Manipulations performed in this study. B. Working model: Nlg3 and DA are part of a pathway responsible for the modulation of social behaviour after isolation in males. Nlg3 would be required downstream of DA, for proper DA signaling in response to the social environment, with some feedback regulation. Females do require Nlg3 for a typical response to isolation, but other neurotransmitters or neuromodulators, responsive to social experience in females, may be interacting with Nlg3. Blue outline: Drosophila brain outline. Created with BioRender.com.

In conclusion, for the first time, we have demonstrated that social space and sociability are recoverable after isolation in a sex-specific manner. We have also shown that DA is required for the recovery from social isolation in a sex-specific manner and that *nlg3* is required for DA levels to respond to the social environment. If those gene by environment by sex interactions are evolutionary conserved, it might help better understanding the molecular basis of the social difficulties encountered by humans with neurodevelopmental disorders in response to changing social environments.

## 6 Author Contributions

Data acquisition and analysis were carried out by RTY, AMS, JMK, and BW-R; conception: RTY and AFS. The first draft was written by RTY and the final draft edited by RTY, AMS, JMK, BW-R, RD and AFS.

## 8 Acknowledgments

We would like to thank Dr. Mark Bernards, Dr. Repon Saha, and Karina Kaberi for their knowledge and expertise obtaining the dopamine level quantification.

## 10 Data Availability Statement

The datasets analyzed for this study can be found in the Dryad repository (https://doi.org/10.5061/dryad.59zw3r2fs).

