## Supplemental material for "Recovery from social isolation requires dopamine in males, but not the autism-related gene *nlg3* in either sex"

| 2 Days Group Housing | | |
|---|---|---|
| | # of Flies Mated | Percent Mated (%) |
| Males | 69/81 | 85.2 |
| Females | 94/103 | 91.2 |
| **4 Days Group Housing** | | |
| | # of Flies Mated | Percent Mated (%) |
| Males | 55/72 | 76.4 |
| Females | 101/112 | 90.2 |

**Supplemental Table 1.** Percent of flies mated with two or four days of group housing before isolation.

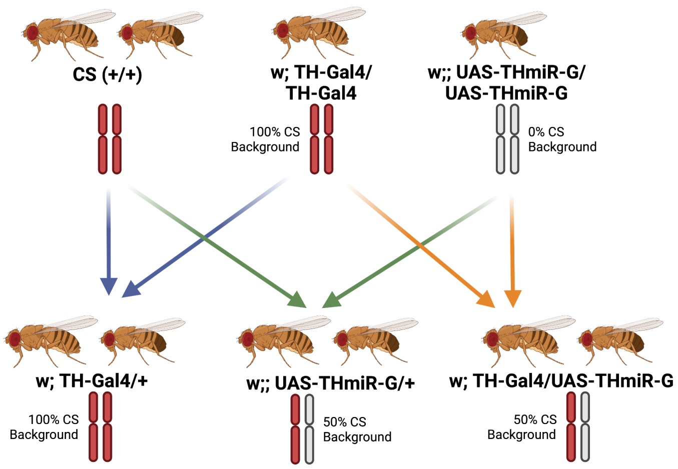


**Supplemental Figure 1.** Schematic of the crosses performed to generate *TH>THmiR-G* and its genetic controls. Created with BioRender.com.

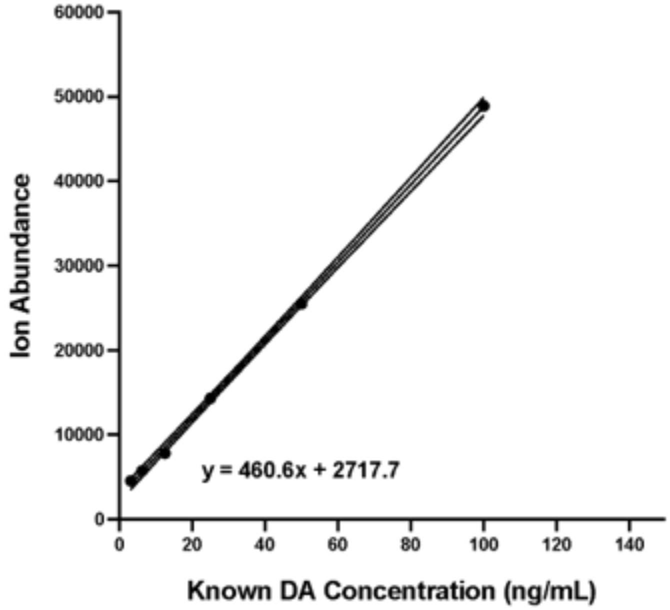


**Supplemental Figure 2.** Standard curves for DA quantification. Equation displayed was used to determine DA concentration. Solid line represents line of best fit. Dotted lines represent SEM.

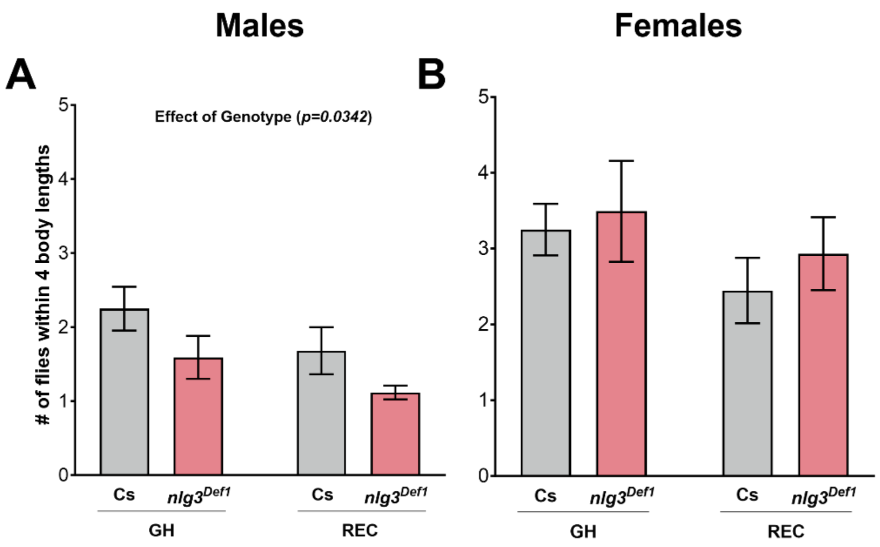


**Supplemental Figure 3.** *nlg3*$^{Def1}$ flies recover after 3 days of group housing following isolation.

**A-B:** Average number of male and female flies within 4 body lengths (± s.e.m) in Cs and *nlg3*$^{Def1}$ flies that were group housed or recovery flies. **A.** Cs and *nlg3*$^{Def1}$ males were not significantly different in number of flies within 4 body lengths in group housed and recovery treatments, but *nlg3*$^{Def1}$ males had fewer flies within 4 body lengths compared to Cs in group housed and recovered treatments (Two-way ANOVA – Effect of Genotype: $F_{1,25} = 5.019$, $p = 0.0342$). **B.** Cs and *nlg3*$^{Def1}$ females also showed no differences in the number of flies within 4 body lengths whether they were group housed or recovery flies (Two-way ANOVA – Effect of Recovery: $F_{1,29} = 1.723$, $p = 0.1996$). n=7-9 with 12–17 flies per trial. GH: Group Housed, REC: Recovery from isolation.
